# Identification of 15 new bypassable essential genes of fission yeast

**DOI:** 10.1101/732438

**Authors:** Aoi Takeda, Shigeaki Saitoh, Hiroyuki Ohkura, Kenneth E. Sawin, Gohta Goshima

## Abstract

Every organism has a different set of genes essential for its viability. This indicates that an organism can become tolerant to the loss of an essential gene under certain circumstances during evolution, via the manifestation of ‘masked’ alternative mechanisms. In our quest to systematically uncover masked mechanisms in eukaryotic cells, we developed an extragenic suppressor screening method using haploid spores deleted of an essential gene in the fission yeast *Schizosaccharomyces pombe*. We screened for the ‘bypass’ suppressors of lethality of 92 randomly selected genes that are essential for viability in standard laboratory culture conditions. Remarkably, extragenic mutations bypassed the essentiality of as many as 20 genes (22%), 15 of which have not been previously reported. Half of the bypass-suppressible genes were involved in mitochondria function; we also identified multiple genes regulating RNA processing. 18 suppressible genes were conserved in the budding yeast *Saccharomyces cerevisiae*, but 13 of them were non-essential in that species. These trends are consistent with a recent independent bypass-of-essentiality (BOE) screening of 142 fission yeast genes conducted with more elaborate methodology (Li *et al*., 2019). Thus, our study reinforces the emerging view that BOE is not a rare event and that each organism may be endowed with secondary or backup mechanisms that can substitute for primary mechanisms in various biological processes. Furthermore, the robustness of our simple spore-based methodology paves the way for genome-scale BOE screening.

## Introduction

A recent genome-wide study using *S. cerevisiae* gave an insight into the ‘evolvability’ of essential cellular processes (Liu *et al*., 2015), which can be also termed ‘bypass-of-essentiality’ (BOE) (Li et al., 2019). The study surveyed the viability of every essential gene disruptant in *S. cerevisiae* (1,106 genes), and found that 9% of the gene disruptants proliferate and form colonies spontaneously (i.e. without artificial mutagenesis). Genome analysis showed that most of the proliferating strains had gained an extra chromosome (i.e. aneuploidy), which is typically an outcome of chromosome missegregation. This is a reasonable path to BOE in *S. cerevisiae*, because its haploid is tolerant to a chromosome gain for 13 of 16 chromosomes (Torres *et al*., 2007). However, we speculated that there might be many more bypassable essential genes in yeast, as some non-bypassable essential gene disruptants might recover their viability by acquiring extragenic mutations, which are rarely introduced without mutagenesis.

Comprehensive identification of suppressor mutations would help to elucidate secondary or backup mechanisms that can substitute for primary mechanisms. Hitherto ‘masked’, these alternative mechanisms may act as the dominant pathways in specific cell types and/or diseased cells. To this end, we designed a BOE screening using the fission yeast *S. pombe*, which has a similar number (1,260) of essential genes to *S. cerevisiae* (Kim *et al*., 2010). A notable difference from *S. cerevisiae* is that *S. pombe* has only 3 chromosomes, and the haploid yeast is inviable when an extra copy of either chromosome I or II (the two larger chromosomes) is inherited (Niwa and Yanagida, 1985). Thus, BOE via extra chromosome gain is likely an infrequent event in *S. pombe*. In the present study, we carried out BOE screening for randomly selected 92 essential genes in *S. pombe*, based on UV mutagenesis of spores in which essential genes were deleted.

## Materials and methods

### Yeast strains and media

A diploid named G29 was used as the host (*h^+^ his2 leu1 ura4-D18 ade6-216* / h^−^ *leul ura4-D18 ade6-210 rpl42.sP56Q*), where *rpl42.sP56Q* allele was used as a counter-selection marker against cycloheximide (Roguev *et al*., 2007). Conventional genetic experiments followed (Moreno *et al*., 1991). Yeast was grown on complete YE5S medium (YE supplemented with leucine, uracil, adenine, lysine, and histidine) or the synthetic PMG or EMM medium at 32°C (plate) or 30°C (liquid). Sporulation was induced on the SPA plate or in the EMMG liquid medium (i.e. PMG containing 1 g/L sodium glutamate instead of 3.75 g/L).

### Gene disruption

Essential genes were selected based on information found in the Pombase database (Wood *et al*., 2012). 92 genes on chromosome II were randomly selected. Conventional one-step replacement was conducted using ~500-bp homologous sequences (5’UTR and 3’UTR of the gene to be deleted) (Krawchuk and Wahls, 1999). A tandem G418-resistance (kanMX) */ura4*+ cassette was used as the selection marker (however, *ura4*+ marker was not actually used for selection). For most genes, we directly generated a linear construct (5’UTR-G418-*ura4*+-3’UTR) by two rounds of PCR using two sets of primers (i.e. nested PCR). In some cases, the PCR fragment was cloned into a vector using an Infusion kit (Takara), and the linear construct was amplified with T7/T3 primer set from the plasmid template. The linear DNA was transformed into the G29 diploid strain using the standard lithium acetate/PEG-mediated method, and disruption of the target gene was confirmed by colony PCR using KOD-Fx-Neo or KOF-ONE kit (Toyobo). When the endogenous gene and G418-*ura4*+ cassette had a similar length, we used a longer version of G418-*ura4*+ cassette to distinguish disrupted and endogenous alleles by length. PCR primers for gene disruptions and their confirmation are listed in Table S1.

### Spore isolation

Exponentially growing heterozygous diploid cells in YE5S (+10 μg/ml G418) were harvested and transferred to EMMG medium (1× 10^6^ cells/ml). After shaking at 200 rpm and 30°C for ≥36 h, cells were harvested, washed twice with sterile water, and resuspended in 0.5% glusulase solution. The solution was shaken at 80 rpm at room temperature overnight to digest non-sporulated cells. The spores were harvested and further treated with 30% ethanol for 30 min (80 rpm, room temperature) to further remove diploid cell contamination. The purified spores were resuspended in sterile water and stored at 4°C.

### Spore quality check

Prior to UV mutagenesis, the viability and purity of spores were determined by plating onto normal YE5S plate and YE5S supplemented with G418 (100 μg/ml) and cycloheximide (100 μg/ml), respectively. No haploid spores were expected to grow on the G418/cycloheximide plate, since an essential gene had been replaced with G418. However, colonies were always formed typically at ~1 × 10^−6^ frequencies. Cells in these colonies were diploids, which we interpreted to be derived from diploid spores generated at low frequency during meiosis; diploid spores would also be resistant to glusulase or ethanol. In cases in which the putative diploid contamination frequency was < 5 × 10^−5^, we moved on to UV mutagenesis and screening. In cases in which the contamination frequency was ≥ 5 × 10^−5^, we discarded the sample and repeated the spore isolation process. The reason for differences in the prevalence of putative diploid spores is unknown.

### BOE screening with UV mutagenesis

1 × 10^7^ spores were plated onto a YE5S plate containing G418 (100 μg/ml) and cycloheximide (100 μg/ml), followed by UV irradiation (90 × 100 μJ/cm^2^: UV Crosslinker, CL-1000, 254 nm, 100 V, 8 W [UVP/Analytik Jena] or Stratalinker UV crosslinker Model 1800 [Stratagene]). Under these conditions, spore viability was approximately 1%. Cycloheximide allows counter-selection against *rpl42*+ gene; in the presence of cycloheximide, haploids possessing the *rpl42.sP56Q* allele can grow but not parental heterozygous diploids (Roguev et al., 2007). Plates were incubated at 32°C for 7 d. In most cases, we detected colonies. To check if each colony represents BOE or diploid contamination, we replica-plated onto EMM minus adenine and SPA plates. After checking spore formation by iodine treatment on SPA, we selected the Ade- and non-spore-forming colonies as candidate BOE haploids; colonies that did not match this criterion were likely diploids and disregarded. The candidate colonies were subjected to colony PCR, with which the disruption of the target gene was reconfirmed (see Fig. 2B). For *top3*Δ, we performed mutagenesis in a *rad13*Δ (DNA repair-deficient) background, in order to decrease UV power (1,500 μJ/cm^2^, 5% viability) and avoid cytotoxicity. However, since we obtained expected BOE results for *cut7*Δ in the presence of *rad13*+, we did not introduce *rad13*Δ for any other genes.

**Figure 1.**
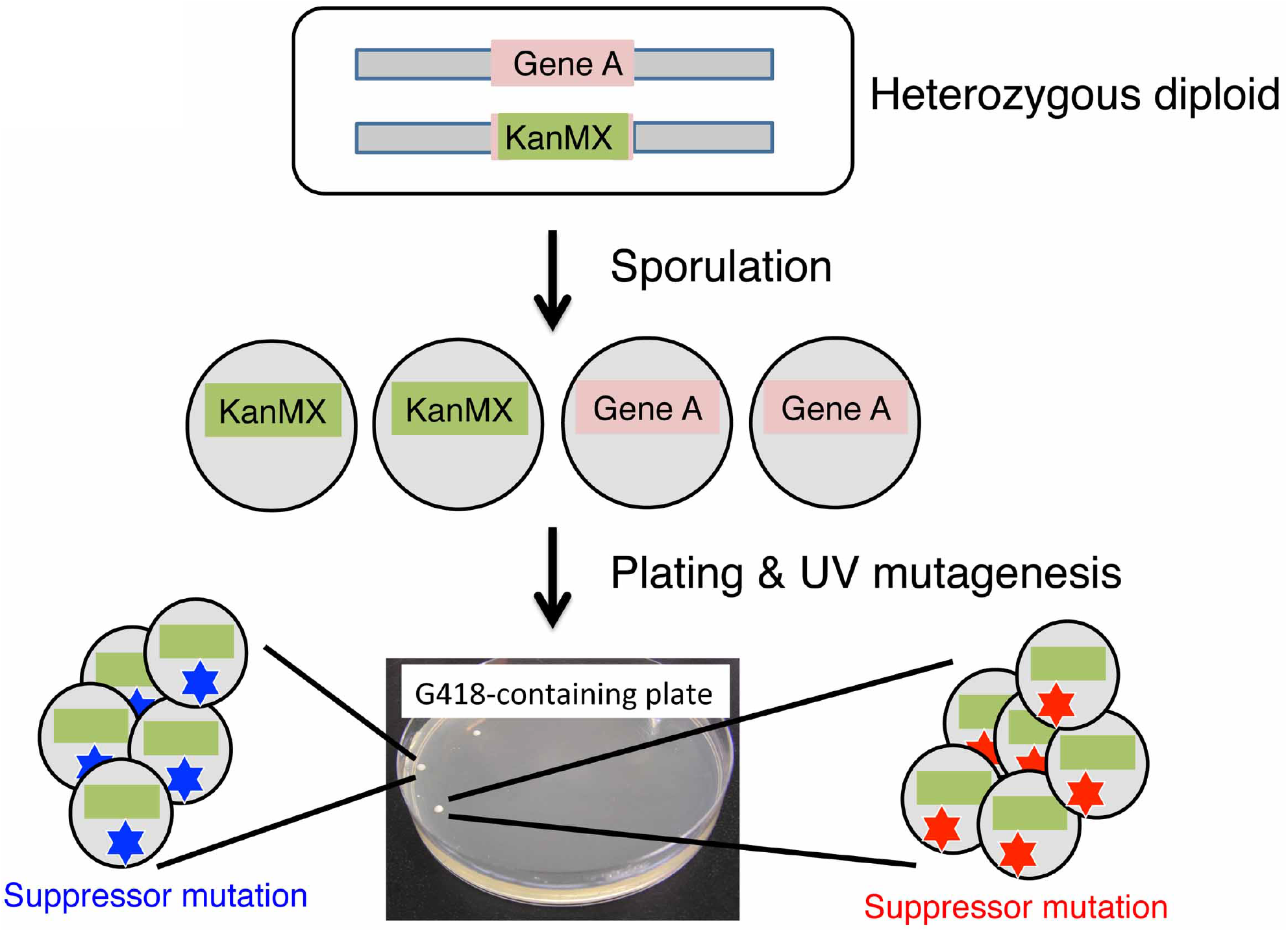
Flowchart of BOE screening using haploid spores of fission yeast. A copy of an essential gene (named ‘A’ in this figure) is replaced by the G418-resistant cassette (KanMX) in the diploid strain. This heterozygous diploid is viable since another copy of gene A remains intact. The diploid is sporulated in the nitrogen-limited medium. Spores with or without gene A are obtained at 1:1 ratio. The spores are spread onto G418-containing plate and then immediately irradiated with UV for mutagenesis. Only a spore with a suppressor mutation can grow and form a colony on the medium. The lack of gene A is confirmed by colony PCR.

**Figure 2.**
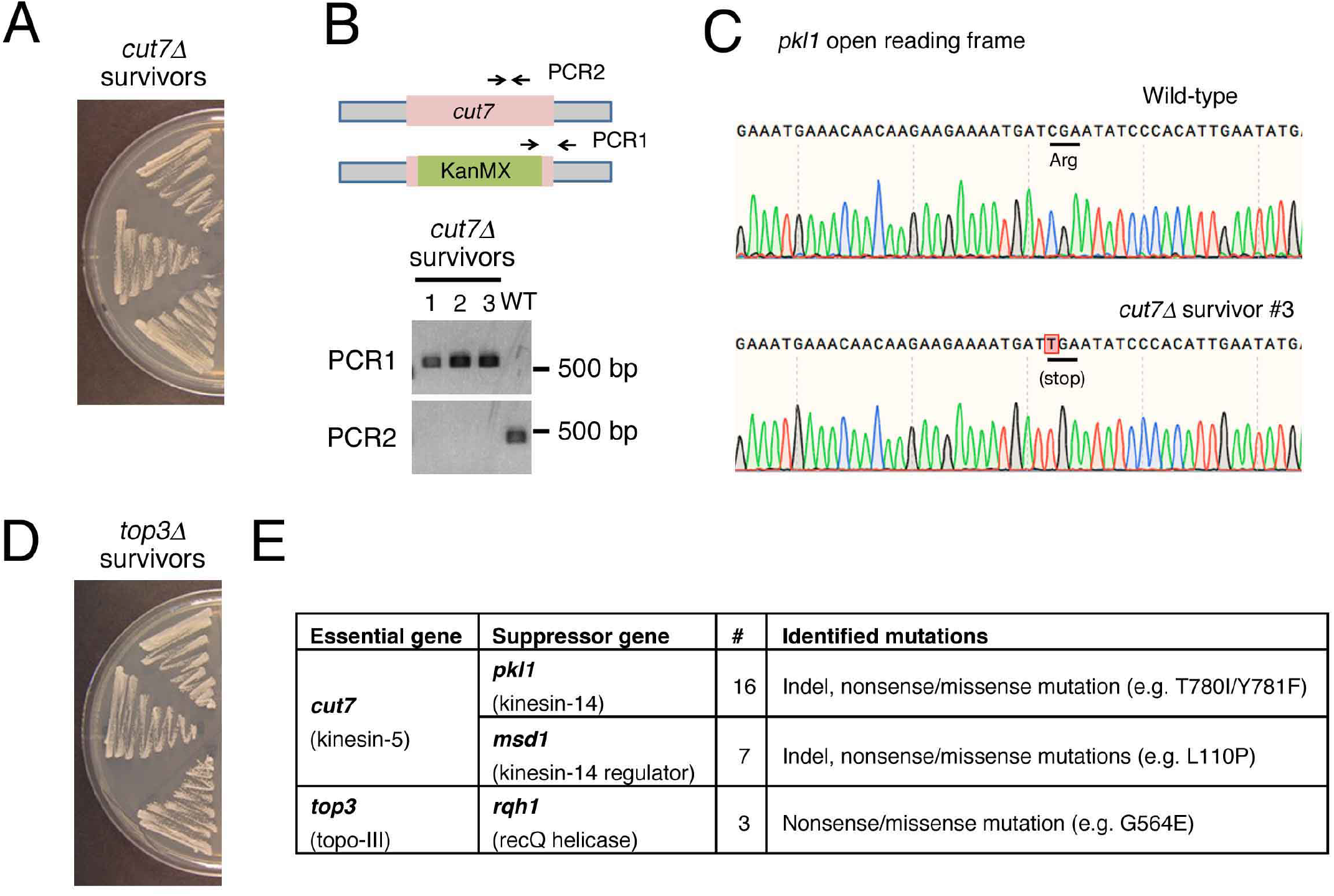
Successful identification of known BOE. (A) Three viable strains after UV mutagenesis of *cut7*Δ strain. In total, we obtained 38 BOE strains after UV mutagenesis of 6 × 10^7^ *cut7*Δ spores. (B) PCR to verify the lack of *cut7* gene for 3 haploid strains that formed colonies. (C) Sequencing result to show the appearance of a premature stop codon in *pkll* gene in a *cut7*Δ BOE strain. (D) 3 viable strains after UV mutagenesis of *top3Δ* spores. (E) Summary of extragenic suppressor mutations of *cut7*Δ and *top3*Δ.

### Whole-genome sequencing and sequence analysis

To identify suppressor mutations, bulk segregant analysis was performed. Survivor strains were crossed with a wild-type strain, and spores were plated on G418-containing YE5S plates. After 7 d, ~1,000 colonies were collected and DNA was extracted with Dr. GenTLE (Takara). Genomic DNA (1 μg) was sequenced by BGI or Novogene (1 Gb), and the reads were mapped to the reference genome (Schizosaccharomyces_pombe.ASM294v2.genebank.gb) using CLC Genomics Workbench. Unique and homogenous Indels and SNPs identified for each strain were manually inspected using Integrative Genomics Viewer (IGV).

## Results & Discussion

Fig. 1 illustrates the scheme of our BOE screening. A heterozygous diploid in which one copy of an essential gene was replaced with a drug (G418)-resistant marker was sporulated. The spores were plated on G418-containing plates and simultaneously mutagenized by UV irradiation. If a haploid colony is obtained on this plate, it has likely acquired a suppressor mutation(s), indicating that the essentiality of the gene has been bypassed.

We first applied this method to two gene disruptants, *cut7*Δ (kinesin-5) and *top3*Δ (type I topoisomerase), the lethality of which is known to be suppressed by the loss of function of Pkl1 (kinesin-14) and Rqh1 (recQ helicase), respectively (Goodwin *et al*., 1999; Olmsted *et al*., 2014; Syrovatkina and Tran, 2015). For *cut7*Δ, we obtained a total of 8 haploid colonies in the first experiment and 30 more in a later experiment, in which 5-fold more spores were mutagenized (Fig. 2A, B). We analysed 26 colonies by target sequencing of *pkl1* and *msd1* genes (Msd1 is a positive regulator of Pkl1 (Yukawa *et al*., 2015)), whole-genome sequencing, and/or genetic linkage test (*pkl1* locus is close to the *rpl42* locus, at which a mutation was introduced to confer cycloheximide resistance in our strain). The combined results suggested that suppressor mutations reside in *pkl1* for 19 strains and in *msd1* for the remaining 7 strains (Fig. 2C, E). Mutagenesis of *top3*Δ yielded 3 haploid strains (Fig. 2D), and direct sequencing of the *rqh1* gene identified a mutation in all cases (Fig. 2E). Thus, our screening successfully elucidated known BOE relationships.

We then expanded the screening to 92 essential genes located on chromosome II. For 20 of these, we obtained 1~17 haploid colonies, which corresponds to 22% (Fig. 3). This frequency is much higher than that obtained in the previous mutagenesis-free screening in *S. cerevisiae* (Liu et al., 2015).

**Figure 3.**
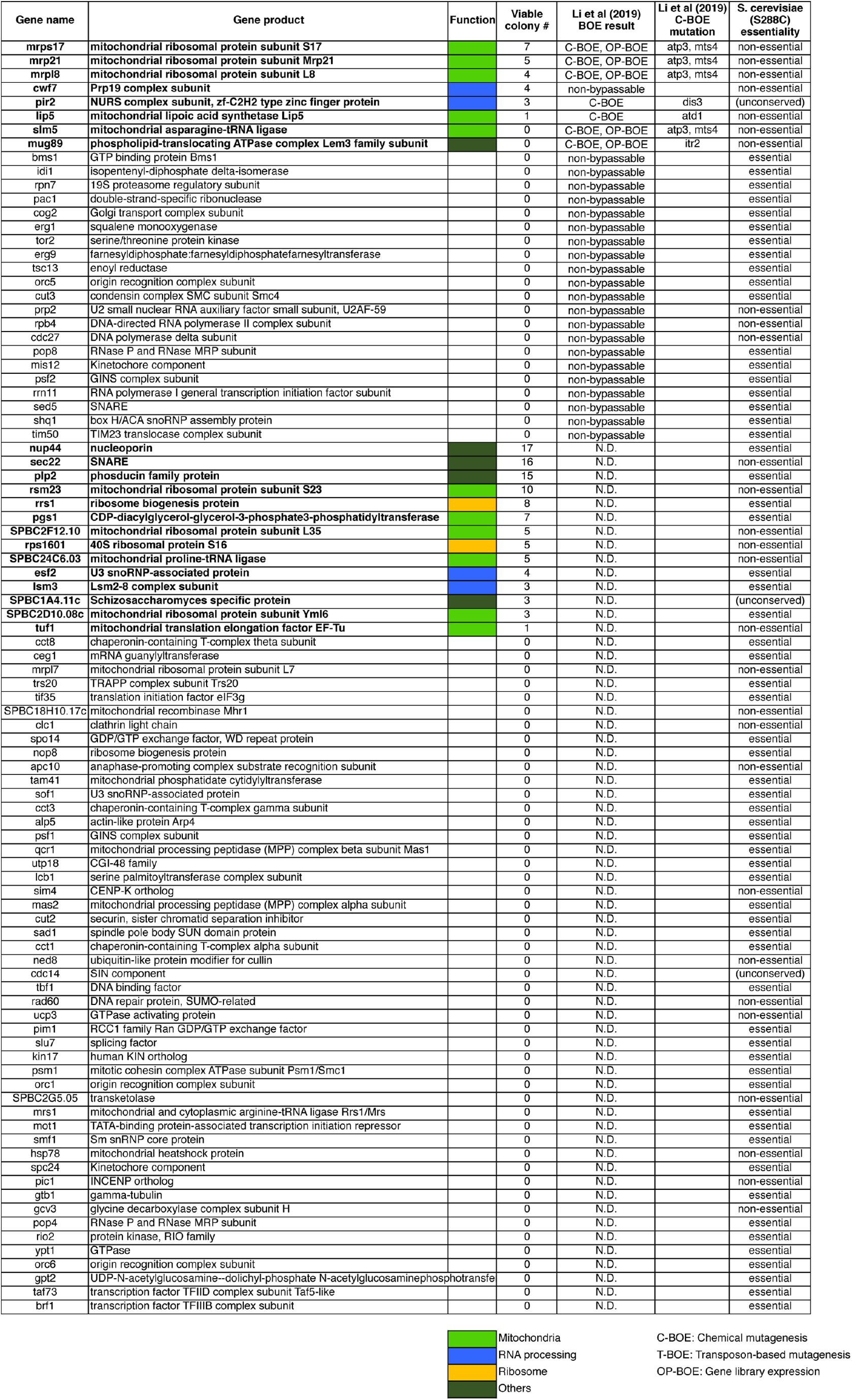
Summary of BOE screening. The gene product was listed based on the information found in Pombase. Information on other BOE screens (Li et al. 2019) and *S. cerevisiae* orthologues (*Saccharomyces* Genome Database (SGD)) are also listed. Bold letters indicate bypassable essential genes identified in our study and/or Li et al. (2019).

The 20 suppressible genes possess divergent known biological functions. 10 genes (50%) were related to mitochondrial function (Fig. 4A). This may be partly explained by the fact that, in the regular medium containing >2% glucose, cell proliferation does not depend much on mitochondrial respiration (Takeda *et al*., 2015). 6 genes were associated with RNA processing and ribosome functions; the basis of these trends are unclear. Overall, 90% of the genes had clear orthologues in *S. cerevisiae* and *H. sapiens*, indicating that BOE is not limited to unconserved genes (Fig. 4B, C). However, the orthologues of 70% genes were reported to be non-essential in *S. cerevisiae* (Fig. 4D). An obvious next step would be to identify suppressor mutations of each survivor to understand how an essential mechanism can be bypassed.

**Figure 4.**
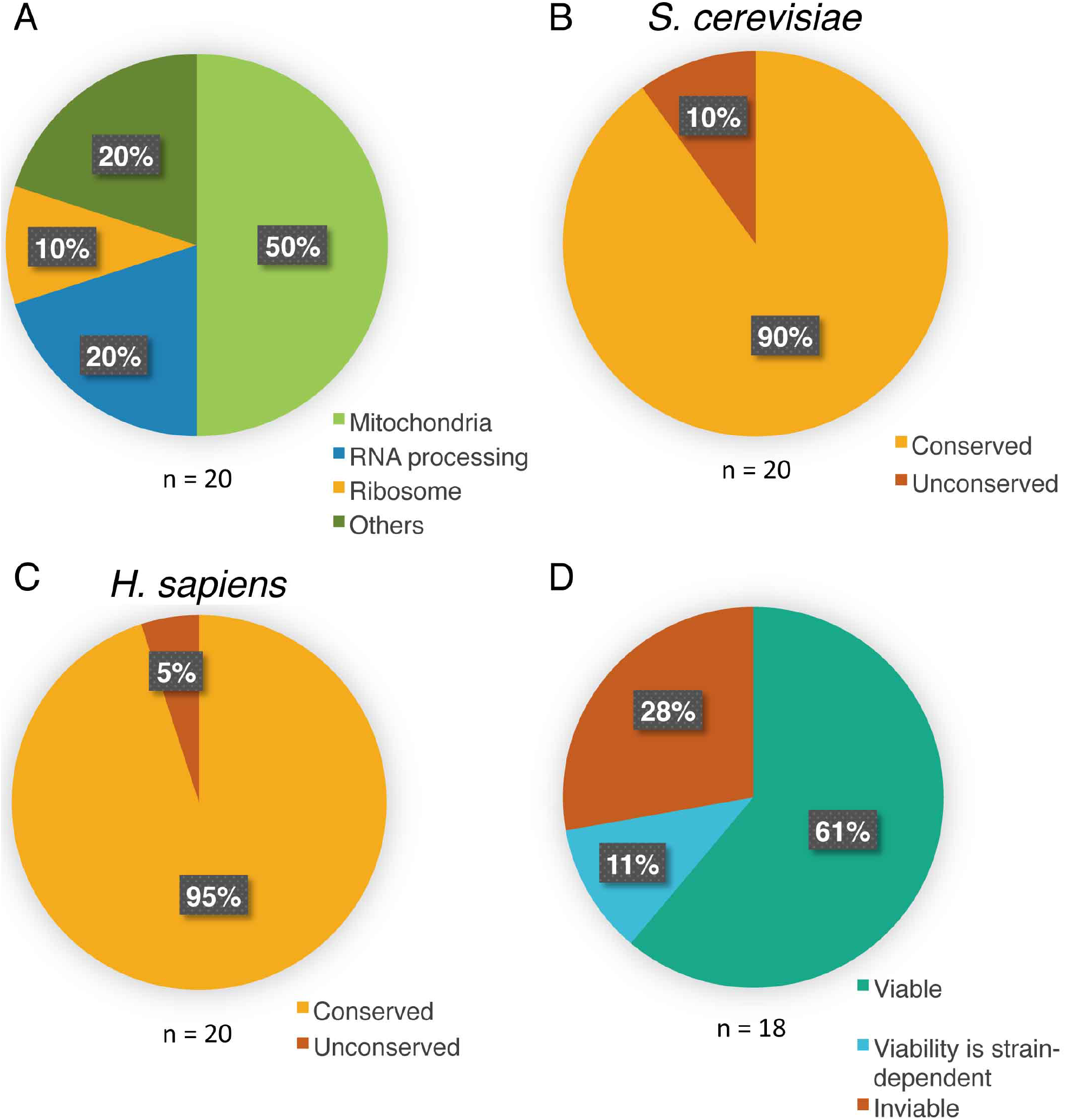
Features of bypassable essential genes. (A) Classification of the function of 20 bypassable essential genes identified in this study. (B, C) Conservation of the identified genes in *S. cerevisiae* (B) or *H. sapiens* (C). (D) Essentiality of the *S. cerevisiae* orthologues (based on *Saccharomyces Genome Database (SGD)*).

In the course of this project, a conceptually identical study was published (Li et al., 2019). In this study using *S. pombe* haploid, BOE was screened by 3 methods: chemical mutagenesis (termed C-BOE), transposon-based mutagenesis (T-BOE), and gene library overexpression (OP-BOE). While C-BOE is the most similar approach to ours, the methodology employed is different. Li et al (2019) did not use spores; instead, the essential gene disruptant was kept viable by the transformation of a plasmid that contains the deleted gene: if a colony that had lost the plasmid was recovered, it was interpreted to indicate BOE. Li et al. (2019) obtained survivors for 27% of the genes in one or more BOE assays, which is a similar frequency to ours.

Coincidentally, in the two studies, 29 common essential genes were screened. Upon comparison, 21 genes were not bypassable in both studies, whereas 5 genes were common BOE hits. 2 and 1 hits were uniquely found in their and our studies, respectively (Fig. 3). Thus, the comparison indicates that both screens had a good agreement in bypassability, but also suggests that a single screen cannot identify all possible BOE.

Li et al (2019) further identified suppressor genes responsible for BOE. For example, they found that the mutation/overexpression of 6 components of the 19S proteasome compensates for mitochondrial dysfunction, suggesting a link between proteasome alteration and mtDNA dispensability. To test if our screen identified the same set of extragenic suppressor genes as Li et al., we determined the whole-genome sequences of 3 BOE strains for *mrpl8* (mitochondrial ribosome protein), which was a C-BOE hit in Li et al (2019). Interestingly, we identified mutations in *atp1* (F1-F0 ATP synthase alpha subunit (Falson *et al*., 1991)) and *rpt3* (19S proteasome base subcomplex ATPase subunit (Kitagawa *et al*., 2014)), which are very similar to what were found in Li et al. (2019) (*atp3;* F1-F0 ATP synthase gamma subunit: *mts4;* 19S proteasome regulatory subunit (Wilkinson *et al*., 1997)). In addition, our screen uniquely identified *hul5* (HECT-type ubiquitin-protein ligase E3 (Fang *et al*., 2011)), which might function upstream of the proteasome. *pir2* (RNA silencing factor (Sugiyama *et al*., 2016)) was another common hit, and Li et al. (2019) reported a single gene mutation in *dis3* (exosome 3’-5’ exoribonuclease subunit (Murakami *et al*., 2007). However, we could not find *dis3* mutations in any of the 3 BOE strains we obtained, indicating that other genes had acquired suppressive mutations.

In summary, we have established an alternative sensitive—and perhaps less labour-intensive—methodology for mutagenesis-based BOE screening in fission yeast, and expanded the list of genes whose essentiality is bypassable. Our methodology allows for a straightforward scale-up of the screen, from which we expect to reveal masked cellular mechanisms.

## Acknowledgements

We are grateful to Ye Dee Tay for various advices on this project; Ulzii Enhjin and Yukina Chiba for primer design; Elsa Tungadi and Juyoung Kim for their help in gene disruption and screening; Kazuma Uesaka and Kunio Ihara for their help in sequence analysis; Iain Hagan, Masamitsu Sato, and National Bio-Resource Project (NBRP) Japan for yeast strains and plasmids. This study originated when G.G. took a sabbatical in the K.E.S. laboratory. Work in the laboratory of K.E.S was supported by the Wellcome Trust (094517, 210659). The subsequent study in the G.G. laboratory was supported by JSPS KAKENHI (19K22383). H.O. laboratory is supported by the Wellcome Trust (098030, 206315).

## Author contributions

G.G. conceived the project. G.G., S.S. H.O. and K.E.S. designed the research. A.T. and G.G. performed experiments. A.T., K.E.S. and G.G. analysed the data. S.S. and K.E.S. contributed resources. G.G. wrote the paper. S.S., H.O. and K.E.S. reviewed and edited the paper.

